## Supplemental Figures and Tables for "Regulation of heterotopic ossification through local inflammatory monocytes in a mouse model of aberrant wound healing"

**Supplemental Figure 1.** Plasma cyto/chemokine levels. Data are shown as the median and interquartile range. Changes in cytokines and chemokines across day 3 and day 7 vs day 0 were analyzed by an analysis of variance (ANOVA) with post-hoc Dunnett test (n=3 mice/time point). Non-heteroscedastic data identified by Levene's test for homogeneity of variances were alternatively analyzed by Welch statistic and post-hoc Dunnett T3. Degrees of freedom (df or df1) across samples = 2. F statistic and significant post-hoc p-values respectively: CXCL1: 0.587, CXCL2: 0.388, CCL2: 1.259, CCL3: 2.295, CCL4: 0.178, G-CSF: 2.736, GM-CSF: 1.099, IL-1b: 6.732, p(D0 vs. D3)=0.032, p(D0 vs. D7)=0.036, IL-6: 11.394, p(D0 vs. D3)=0.009,TNF-α: 0.303, TGF-β1: 2.746, p(D0 vs. D3)=0.009,TGF-β2: 12.294, p(D0 vs. D3)=0.027, p(D0 vs. D7)=0.007, TGF-β3: 0.303, LIF: .994. *p<.05 **p<.01.

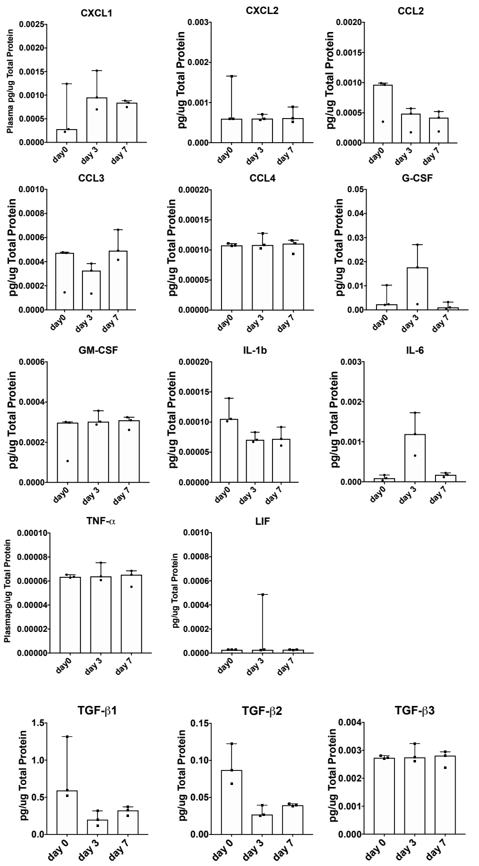


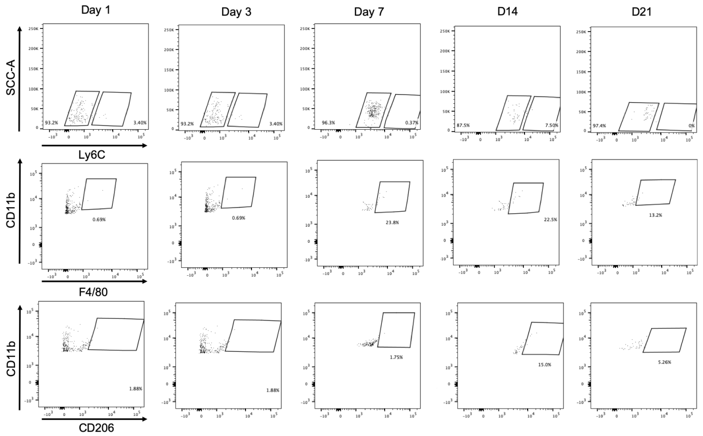


**Supplemental Figure 2.** Negative gates of HO site flow cytometry. Flow cytometry plots showing negative gating strategy for Ly6C, F4/80 and CD206 across the individual time points.

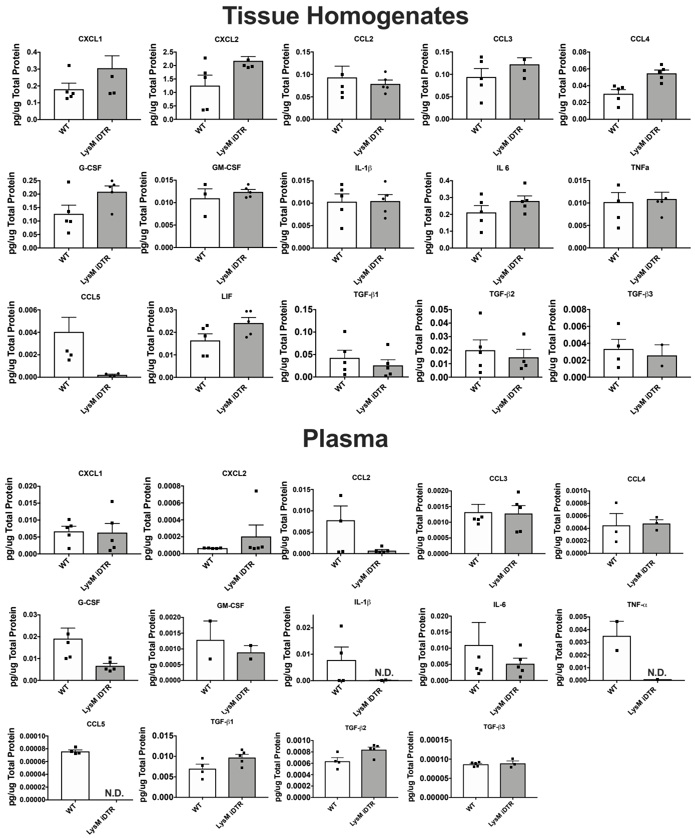


**Supplemental Figure 3.** Homogenate and Plasma levels of cytokines/chemokines after monocyte depletion using LysMCre-*iDTR* or control WT mice (n=5 mice/group). Injury site homogenate and plasma at day 3 from LysMCre-iDTR and liter mate control mice, where monocytes/macrophages were depleted by pre-injection of diptheria toxin (DT) two days before the B/T, the day of B/T and at day 2 after the B/T.

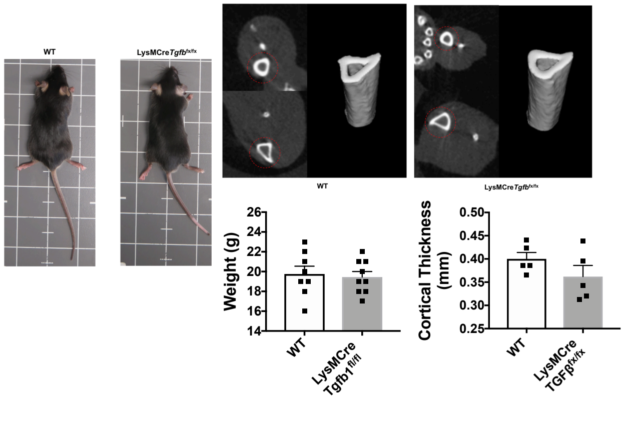


**Supplemental Figure 4.** LysMcre/Tgfb1$^{fl/fl}$ and wild type mice do not differ in size, weight, or tibial thickness. A. Top: images of wild type and LysMcre/Tgfb1$^{fl/fl}$ mice. Bottom: mean weights from mice in either WT or LysMcre/Tgfb1$^{fl/fl}$ groups. B. Quantification of mean cortical tibial thickness from MicroCT scans of WT and LysMcre/Tgfb1$^{fl/fl}$ mice (n=5/group, t=-0.121, df=8, p=0.907)

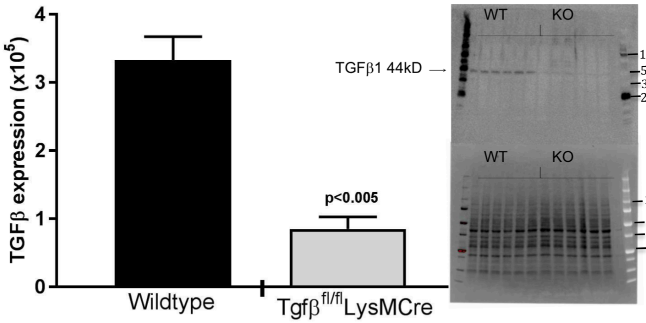


**Supplemental Figure 5**. TGF-β1 expression is reduced in LysMCre-*Tgfb1*$^{fl/fl}$ bone marrow derived macrophages. Bone marrow was flushed from 4-week-old WT (C57B6) or KO (LysMCre-*Tgfb1*$^{fl/fl}$) mice and macrophages induced in culture with 30ng/ml M-CSF for 5 days. Western blot revealed a significant decrease in TGF-β1. Graph represents TGF-β1 expression (seen in Western blot; right, top panel) normalized to loading signal (right, bottom panel). n=3/gp (run in duplicate; t test: p=0.0004, t=11.15, df=4) error bars represent SD.

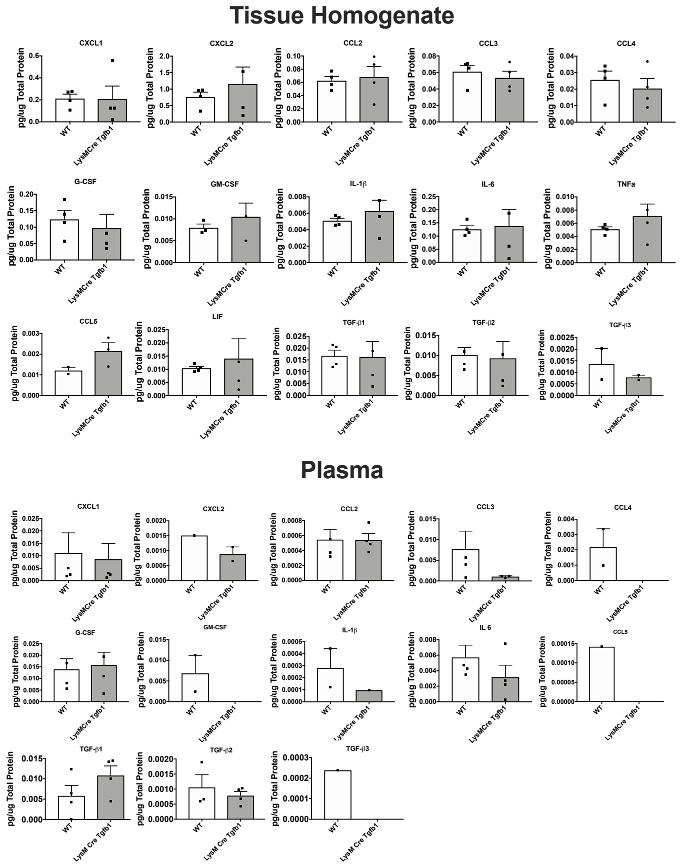


**Supplemental Figure 6.** Homogenate and Plasma levels of cytokines/chemokines after monocyte depletion using LysMCre-*Tgfb1*$^{fl/fl}$ mice (n=4 mice/group).

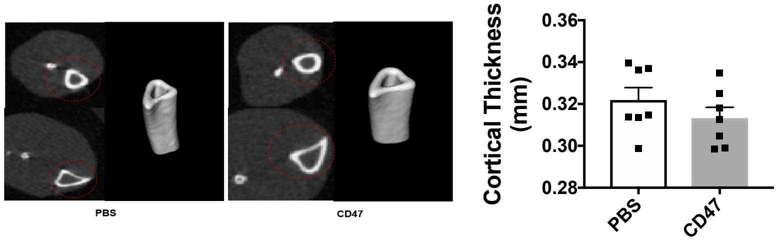


**Supplemental Figure 7.** Treatment with CD47 activating peptide p7N3 does not result in changes in tibial thickness of mice. Top: Quantification of mean cortical tibial thickness from MicroCT scans (n=7/group, t=1.103, df=12, p=0.292). Bottom: Representative MicroCT scan images from tibias of PBS and CD47 peptide (p7N3) treated mice.

**Supplemental Table 1**: Top gene expression of day 3 TGF-β expressing clusters.

| p_val | avg_logFC | pct.1 | pct.2 | p_val_adj | cluster | gene |
|---|---|---|---|---|---|---|
| **0** | 1.47036156 | 0.979 | 0.66 | 0 | 0 | Plac8 |
| **0** | 1.32737263 | 0.907 | 0.353 | 0 | 0 | Ifitm6 |
| **3.73E-179** | 1.31302633 | 0.817 | 0.512 | 6.57E-175 | 0 | Chil3 |
| **4.79E-299** | 1.15385338 | 0.848 | 0.343 | 8.43E-295 | 0 | Ly6c2 |
| **1.97E-266** | 1.07680069 | 0.98 | 0.844 | 3.47E-262 | 0 | Thbs1 |
| **0** | 1.06899311 | 0.983 | 0.68 | 0 | 0 | Hp |
| **0** | 1.02867456 | 0.967 | 0.623 | 0 | 0 | Gsr |
| **0** | 0.98573785 | 0.825 | 0.316 | 0 | 0 | Vcan |
| **2.27E-288** | 0.94741447 | 0.832 | 0.315 | 3.99E-284 | 0 | F10 |
| **4.51E-283** | 0.94567273 | 0.77 | 0.221 | 7.94E-279 | 0 | Gm9733 |
| **0** | 2.4728712 | 0.84 | 0.292 | 0 | 1 | Arg1 |
| **5.01E-158** | 2.05073848 | 0.97 | 0.884 | 8.81E-154 | 1 | Spp1 |
| **3.25E-66** | 1.63075966 | 0.512 | 0.29 | 5.72E-62 | 1 | Cxcl3 |
| **5.61E-37** | 1.12059119 | 0.165 | 0.048 | 9.87E-33 | 1 | Mmp12 |
| **5.00E-108** | 1.06784211 | 0.988 | 0.947 | 8.81E-104 | 1 | Pf4 |
| **3.52E-133** | 1.05879712 | 0.989 | 0.936 | 6.20E-129 | 1 | Ctsl |
| **2.60E-160** | 1.02772177 | 0.865 | 0.568 | 4.58E-156 | 1 | Hilpda |
| **4.69E-17** | 1.02137961 | 0.228 | 0.128 | 8.25E-13 | 1 | Ppbp |
| **4.31E-81** | 0.9504383 | 0.49 | 0.215 | 7.59E-77 | 1 | Cd36 |
| **3.46E-178** | 0.9223303 | 0.988 | 0.899 | 6.10E-174 | 1 | Mif |
| **1.72E-86** | 0.77324378 | 0.933 | 0.832 | 3.03E-82 | 3 | H2-Aa |
| **3.61E-111** | 0.76874144 | 0.997 | 0.974 | 6.36E-107 | 3 | Cd74 |
| **3.40E-78** | 0.72232396 | 0.906 | 0.748 | 5.99E-74 | 3 | H2-Eb1 |
| **1.79E-149** | 0.67948462 | 0.991 | 0.816 | 3.14E-145 | 3 | Aif1 |
| **4.87E-164** | 0.66808656 | 1 | 0.921 | 8.57E-160 | 3 | Ctsc |
| **3.51E-87** | 0.6639899 | 0.96 | 0.898 | 6.18E-83 | 3 | H2-Ab1 |
| **3.39E-86** | 0.59492665 | 0.978 | 0.911 | 5.98E-82 | 3 | Marcksl1 |
| **4.67E-123** | 0.57249586 | 0.974 | 0.758 | 8.22E-119 | 3 | AF251705 |
| **4.22E-79** | 0.52282696 | 0.993 | 0.949 | 7.44E-75 | 3 | Tgfbi |
| **1.81E-88** | 0.51670001 | 0.979 | 0.716 | 3.19E-84 | 3 | Ccr2 |

**Supplemental Table 2:** Common Macrophage Subsets at HO site in PBS and CD47 activating peptide treatment.

| Cluster # | | Gene |
|---|---|---|
| **PBS** | **CD47** | |
| 4 | 1 | Spp1 |
| | | Arg1 |
| | | Lgals3 |
| | | Cstb |
| | | Cd36 |
| | | Prdx1 |
| | | Mmp12 |
| 3 | 3 | Plac8 |
| | | Ccr2 |
| | | Cd52 |
| | | Il1b |
| | | H2-Eb1 |
| | | H2-Aa |
| 1 | 2 | Sepp1 |
| | | C1qa |
| | | Apoe |
| | | C1qb |
| | | C1qc |
| | | Aif1 |
| | | Trem2 |
| | | Lyz2 |

**Supplemental Table 3**: Unique Macrophage Subsets at HO site with CD47 activating peptide treatment

| Gene | Avg. Fold Change | % Expression in cluster | % Expression all other clusters | Adjusted p-val | Cluster # |
|---|---|---|---|---|---|
| **PBS** | | | | | |
| Ccl4 | 1.92741015 | 0.594 | 0.175 | 1.89E-171 | 9 |
| Ccl3 | 1.428509938 | 0.544 | 0.125 | 4.42E-194 | 9 |
| Mrc1 | 1.325374474 | 0.847 | 0.249 | 9.71E-263 | 9 |
| Cd83 | 1.305317037 | 0.797 | 0.189 | 1.29E-302 | 9 |
| Isg15 | 2.096509936 | 0.908 | 0.2 | 5.41E-177 | 12 |
| Ccl12 | 1.981830311 | 0.638 | 0.093 | 6.41E-153 | 12 |
| Ifit3 | 1.893926081 | 0.85 | 0.058 | 0 | 12 |
| Irf7 | 1.803168506 | 0.932 | 0.187 | 1.11E-197 | 12 |
| Fcgr1 | 1.502130958 | 0.937 | 0.243 | 6.60E-152 | 12 |
| Ifit1 | 1.44619128 | 0.643 | 0.059 | 4.30E-247 | 12 |
| **CD47** | | | | | |
| Folr2 | 1.688539551 | 0.671 | 0.099 | 0 | 6 |
| Mrc1 | 1.623389766 | 0.925 | 0.243 | 0 | 6 |
| Ccl12 | 1.615203405 | 0.58 | 0.077 | 0 | 6 |
| Cbr2 | 1.444270202 | 0.629 | 0.071 | 0 | 6 |
| F13a1 | 1.294819395 | 0.765 | 0.203 | 0 | 6 |
| Fcrls | 1.281069482 | 0.699 | 0.092 | 0 | 6 |
| Clec10a | 1.234401091 | 0.603 | 0.126 | 2.6929E-301 | 6 |
| Cxcl3 | 4.773273925 | 0.963 | 0.1 | 4.6591E-167 | 12 |
| Ccl3 | 4.597986709 | 1 | 0.22 | 3.71915E-95 | 12 |
| Csf3 | 3.566634486 | 0.768 | 0.022 | 0 | 12 |
| Serpinb2 | 3.413322798 | 0.707 | 0.011 | 0 | 12 |
| Il1a | 3.274139858 | 0.976 | 0.022 | 0 | 12 |
| Arg1 | 2.644316173 | 0.939 | 0.114 | 1.8472E-131 | 12 |
| Il1rn | 2.623427277 | 0.915 | 0.218 | 7.91127E-72 | 12 |
| Nos2 | 2.210727868 | 0.78 | 0.01 | 0 | 12 |
| Inhba | 2.065551886 | 0.805 | 0.063 | 1.3879E-168 | 12 |
| Egln3 | 1.497285254 | 0.72 | 0.022 | 0 | 12 |
| Ero1l | 1.483284038 | 0.841 | 0.19 | 1.16011E-66 | 12 |

**Supplemental Table 4:** Taqman primer/probe assays used for QPCR.

| Gene | | Lot | Assay ID |
|---|---|---|---|
| *Gapdh* | FAM | 1593842 | Mm99999915_g1 |
| *Nos1* | FAM | 1732158 | Mm01208059_m1 |
| *Arg1* | FAM | 1712842 | Mm00475988_m1 |
| *Mrc1* | FAM | 1721427 | Mm01329362_m1 |
| *Tgfβ1* | FAM | 1730281 | Mm01178820_m1 |
